# QuadCleave: Enzymatic cleavage and analysis of DNA G-quadruplexes

**DOI:** 10.64898/2026.05.15.725486

**Authors:** Rachael Baker, Alaina Hrysikos, Olivia Lewis, Randi Dias, Emma Younger, Tarah Anasseri, Diti Patel, Samuel Leiszler, Christopher Syed, Krishna Patel, Chirag Lodha, Levi Diggins, Daniel Ross, Shane Donahue, Olivia McLean, Shadika Panta, Camila Pacocha, Matthew Gevaert, David Eagerton, Kaushlendra Tripathi, Bidyut K. Mohanty

## Abstract

Prediction of G-quadruplexes (G4), noncanonical DNA secondary structures containing guanine tetrads, can have profound relevance across cancer, neurodegenerative and other genetic diseases, but require experimental validation by instrument intensive methods including CD spectroscopy, fluorescence, NMR, or crystallography. We report a novel, inexpensive, efficient and scalable 2-enzyme system for confirmation of G4 formation and copy number repeat count. First, QuadCleave, in which Mismatch Endonuclease I was used to repeatably, site-specifically and selectively cleave oligodeoxynucleotides containing exclusively G4-forming sequences. Second, HaeIII enzyme identified G4-forming sequences that can also generate GG/CC base pairs, and a sequence’s cleavage by both established it in C9ORF72, the most mutated gene in amyotrophic lateral sclerosis (ALS) and frontotemporal dementia (FTD). These findings establish ubiquitously accessible experimental validation of G4 forming sequences and their copy number repeat assessment, with specificity to C9ORF72 oligodeoxynucleotides whose repeat expansion has critical clinical implications in a subpopulation of ALS and FTD patients.

## Introduction

Interspersed in the B-form DNA of the human genome are various noncanonical structures including G4s, i-motifs, hairpins, cruciform structures, human fragile sites containing minisatellites, and triple helices formed by homopurine-homopyrimidine trinucleotide repeats^1^. G4s (**Fig. 1 A-C**) are four-stranded DNA secondary structures that contain two or more stacks of guanine tetrads with each tetrad holding four guanines joined by Hoogsteen base association in a plane^2^. A G4 can be intermolecular containing the four strands of DNA (**Fig. 1 A and B**) or an intramolecular G4 (**Fig. 1 C**). Further, a G4 can be parallel with all four strands in one direction (**Fig. 1 A**), antiparallel with two strands in opposite direction to the other two strands (**Fig. 1 B**) or hybrid with three strands in opposite direction to the fourth strand^3^. We have grouped antiparallel and hybrid G4s into nonparallel G4s^3^. In contrast to G4s, i-motifs^4^ are noncanonical four-stranded DNA structures in which cytosines are held together by hemi-protonated and intercalated cytosine base pairs (C:C^+^ or C:C)^5,6^. The human genome contains thousands of G4-forming^7^ and i-motif-forming sequences^8,9^. A polyguanine-rich DNA sequence and its complementary polycytosine-rich DNA can form G4 and i-motif across each other; however, the two forms can coexist or are mutually exclusive^10^. Several G4-forming sequences are found at the promoter-proximal regions of various oncogenes^11^, including cMyc^12^, BCL2^13^, hRAS^14,15^, HIF1-α^16^, EGFR^17^, PDGF^18^, and VEGF^19,20^ and are implicated in gene expression. The telomeric sequence 5’-GGGTTA-3’ can also form a G4^21,22^, which has been implicated in cancer^23,24^. G4/i-motif-forming sequences are also present in several genes associated with various neurological disorders including Alzheimer’s disease (AD)^25^, fragile X syndrome (FXS)^26^, Parkinson’s disease (PD)^27^, amyotrophic lateral sclerosis (ALS), and frontotemporal dementia (FTD)^28,29^. The most common mutation associated with ALS and FTD is a hexanucleotide repeat expansion (HRE) in intron 1 of the *C9orf72* gene. The hexanucleotide sequence 5’-GGGGCC-3’, which expands from normally present 2-23 copies to 700-1600 copies in certain ALS/FTD patients, forms G4 whereas its complementary sequence 5’-CCCCGG-3’ can form i-motif. Although G4s and i-motifs can be beneficial to the cell, they can adversely affect important genome integrity processes including DNA replication^30,31^, recombination^32^, transcription^33^, and translation^34^ and thus are being studied extensively for the diagnosis and therapeutics of ALS and cancer.

**Fig. 1.**
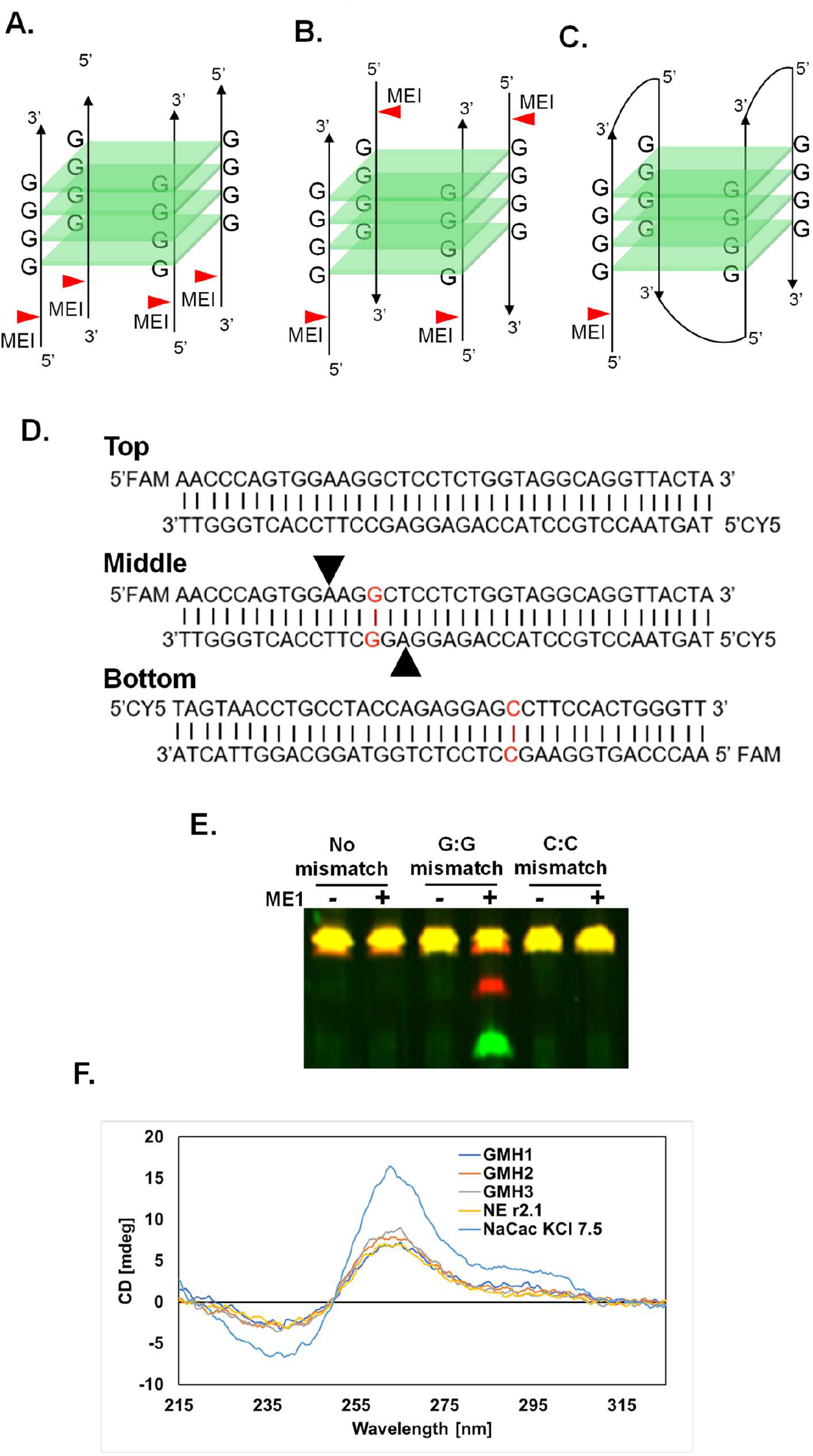
G-quadruplex model and proposed mechanism of Mismatch Endonuclease I (MEI) cleavage G:G mismatches. **A-C**. models of G-quadruplex (G4); **A**. Intermolecular parallel G4, **B**. Intermolecular antiparallel G4, and **C**. Intramolecular antiparallel G4. **D**. Sequences of three FAM- and Cy5-tagged double-stranded DNA used for MEI digestion. **D Top**: DNA with no mismatch; **D Middle**: DNA with a G:G mismatch; **D Bottom**: DNA with a C:C mismatch. **E**. 7M urea-20% polyacrylamide gel showing MEI digestion of FAM- and Cy5-tagged double-stranded DNA. Yellow fluorescence in all lanes is by two intact DNA fragments in the lanes with no enzyme or undigested DNA. The two DNA fragments are tagged with Cy5 (red fluorescence) and FAM (green fluorescence). The two lanes showing “**No mismatch**”, the two lanes showing “**G:G mismatch**” and the two lanes showing “**C:C mismatch**” each contained DNA without (-) and with (+) MEI. In the lane containing DNA with “G:G mismatch” whereas the yellow fluorescence represents the two uncut (Cy5- and FAM-tagged) DNA samples together, the red and green bands represent the truncated Cy5- and FAM-tagged DNA. **F**. CD spectroscopic analysis of a G4-forming 36-nucleotide oligo (GGGGCC)X6 in five different buffers.

A given strand’s potential to form a G4 structure does not necessarily map to actual G4 formation; the latter must be validated experimentally. Various methods are employed to determine the formation of a G4 or an i-motif by a DNA fragment^35^ or its mutants. The common *in vitro* biophysical methods include circular dichroism (CD) spectroscopy, UV-visible absorbance spectroscopy, nuclear magnetic resonance (NMR), gel electrophoresis, size-exclusion high pressure liquid chromatography (SE-HPLC), analytical ultracentrifugation (AUC), and Forster Resonance Energy Transfer (FRET) ^3,11,35^. In addition, G4- and i-motif-antibodies are commercially available to purify and visualize the two groups of DNA structures^36–39^. However, no enzymatic method has yet been developed to study G4s. Most of the biophysical techniques use expensive instruments, which may not be available or accessible to researchers easily. Here we show that Mismatch Endonuclease I (MEI) recognizes and cleaves G4-forming single-stranded DNA (ssDNA) oligodeoxynucleotides (oligos) in a site-specific manner (QuadCleave). MEI is a commercially available (New England Biolabs; NEB) recombinant enzyme previously known to recognize and cleave double-stranded DNA (dsDNA) containing G:G, G:T and T:T mismatches two nucleotides upstream of the mismatch in each strand of the dsDNA. Heretofore unknown for this application, we tested MEI for the cleavage of several single-stranded oligos containing G4-forming- and i-motif-forming sequences of *C9orf72* intron 1, oncogene promoter-proximal regions, and telomeres. We also tested ssDNA oligos that generate other structures or remain linear. We demonstrated MEI’s generation of specific cleavage products of only G4-forming oligos, but not other oligos in various buffers containing components that favor G4 formation and MEI cleavage. Specifically, monovalent cations of potassium and sodium which favor G4 formation, and magnesium which is necessary for MEI activity, were used for the enzymatic activity. The enzyme did not cleave any i-motif-forming ssDNA molecules tested or any other non-G4 forming DNA samples. Our QuadCleave method is reproducible and inexpensive and can thus be efficiently used prior to biophysical analysis. It holds significant promise for G4 analysis and diagnosis in the laboratory. In addition, we have discovered that HaeIII restriction endonuclease enzyme can additionally be used to determine GG/CC base-pairing in oligos containing G4-forming 5’-GGGGCC-3’ sequences and i-motif-forming 5’-CCCCGG-3’ sequences. QuadCleave, in combination with HaeIII, can be used to determine complex structures formed by GGGGCC repeats of the *C9orf72* gene.

### Mismatch Endonuclease I cleaves double-stranded DNA containing G:G mismatch

Since MEI is known to cleave DNA containing a G:G mismatch, we first tested the enzyme for the cleavage of a double-stranded DNA (dsDNA) containing a G:G mismatch. For this, we prepared three dsDNA substrates each containing a FAM-tagged ssDNA oligo and a Cy5-tagged complementary ssDNA oligo (**Fig. 1D**). The first dsDNA substrate did not contain any mismatch (**Fig. 1D Top**); the second substrate contained a G:G mismatch (**Fig. 1D middle**); and the third substrate contained a C:C mismatch (**Fig. 1D bottom**). The substrates were incubated with MEI, and the products were analyzed in a denaturing urea-PAGE gel. Since each dsDNA contains two oligos of the same size, a FAM-tagged oligo (green) and a Cy5 tag oligo (red), the two oligos (green plus red) fractionated at the same position in a denaturing gel and displayed yellow color (**Fig. 1E, all three –MEI lanes**). MEI treatment of the dsDNA without any mismatch did not show any change from undigested DNA (yellow; **Fig. 1E**, with no mismatch, **lanes –MEI and +MEI**). However, MEI treatment of a substrate containing the G:G mismatch showed three bands, a yellow band representing the undigested DNA, a red band and a green band; the latter two representing two cleaved products of the Cy5-tagged and FAM-tagged strands, respectively (**Fig. 1E**, with G:G mismatch shown in red, compare **lanes –MEI and +MEI**). The enzyme cleaved each of the two oligos due to the G:G mismatch. In contrast to the G:G mismatch, MEI did not cleave the DNA substrate containing a C:C mismatch displaying only the yellow band in undigested and digested lanes (**Fig. 1E**, with C:C mismatch shown in red, **lanes –MEI and +MEI**). These results show that MEI recognized and cleaved dsDNA containing the G:G mismatch but did not cleave DNAs without any mismatch or the C:C mismatch.

### Mismatch Endonuclease I cleaves G4-forming hexanucleotide DNA repeats of *C9orf72* gene

To determine if MEI recognizes and cleaves G4-forming DNA oligos containing 5’-GGGGCC-3’ repeats, we designed five FAM-tagged oligos (G-1 - G-5) containing 1-5 repeats of 5’-GGGGCC-3’ sequence flanked by 10 nucleotides on either end (**Supplementary Materials; Table S1**). We also designed five similar FAM-tagged oligos (G-1 NF - G-5 NF) containing the same 1-5 repeats of 5’-GGGGCC-3’ without any flanking sequences (**Supplementary Materials; Table S1**). To design buffers that favor both G4 formation and MEI cleavage, we considered three ions: potassium, sodium, and magnesium. Recently, by using CD spectroscopic analyses, we have tested the role of pH, temperature, and hexanucleotide repeat numbers on G4 formation by oligos containing 5’-GGGGCC-3’ repeats of *C9orf72*^3,11^ and showed that G4 formation by the repeats was favored in a buffer containing 10 mM sodium cacodylate and 100 mM KCl at pH 7.4 (**Fig. 1F**). G4 formation can also occur in the presence of sodium ions replacing potassium ions, although KCl is used most frequently^11^. For MEI digestion, the commercially available buffer contains Tris, NaCl and MgCl_2_. We used four different buffers for MEI cleavage: GMH1 (containing 10 mM sodium cacodylate and 100 mM KCl; pH 7.4), GMH2 buffer (containing 10 mM sodium cacodylate and 50 mM NaCl; pH 7.4), GMH3 buffer (containing 10 mM Tris.HCl and 50 mM NaCl; pH 7.5), and NEBuffer r2.1 (Tris.HCl, 50 NaCl, 10 mM MgCl_2_, pH 7.9). Since G4 formation requires either K^+^ or Na^+^ ions and MEI activity requires Mg_2_^+^ ions, all four buffers contain MgCl_2_ and KCl or NaCl. Denaturing gel analyses of the MEI-treated oligos show that MEI cleaved all five *C9orf72* oligos tested that contain flanking sequences (G1 – G5) generating fragments of specific sizes (**Fig. 2 A-D**). As the number of repeats increased, so did the number of cleaved fragments, which displayed the generation of a specific pattern for each oligo. The generation of the unique binding patterns suggests site-specific cleavage of the G4-forming DNA oligos. To explore the location of cleavage relative to G:G:G:G mismatches, we designed G1-G5 oligos with 10 nucleotides on each side of the G4-forming sequences and demonstrated consistency with MEI’s known digestion of G4-forming oligos upstream of G:G:G:G mismatches.

**Fig. 2.**
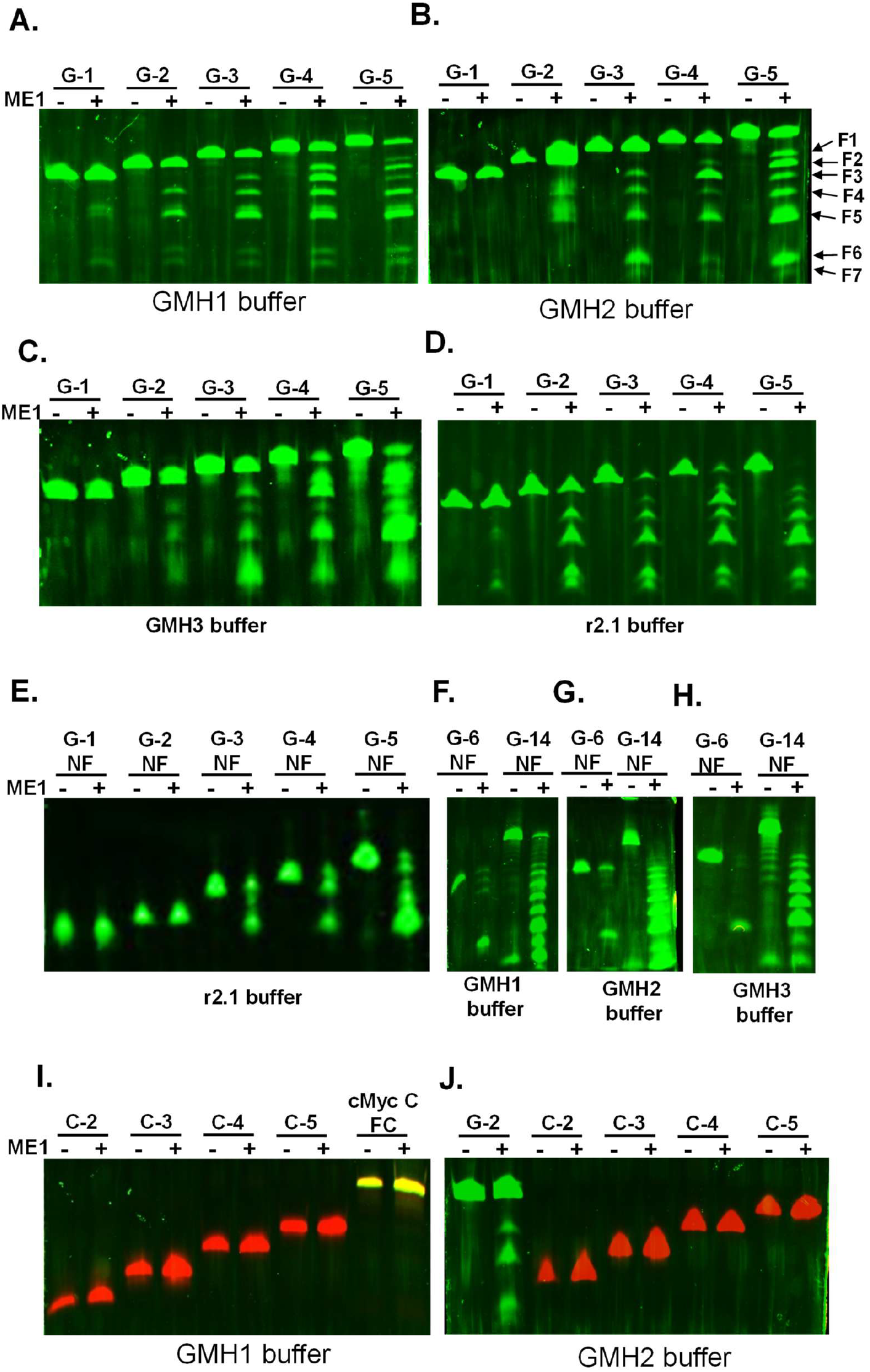
7M urea-20% polyacrylamide gels show that Mismatch Endonuclease I (MEI) cleaves polyguanine-rich DNA oligos, but not polycytosine-rich DNA oligos of *C9orf72* in a repeat-dependent manner. Lanes MEI (-) and (+) in all gels represent DNA samples without and with MEI treatment. **A-D**. G-1 to G-5 oligos (containing 1-5 repeats of 5’-GGGGCC-3’ and flanking sequences) were treated with MEI in GMH1, GMH2, GMH3 and NEBuffer r2.1. **E**. G-1 NF to G-5 NF oligos (with 1-5 repeats of 5’-GGGGCC-3’ sequence) without or with MEI in NEBuffer r2.1. **F-H**. G-6 NF and G-14 NF (with 6 and 14 repeats of 5’-GGGGCC-3’ sequence) without and with MEI in GMH1, GMH2 and GMH3 buffers respectively. **I**. Cy5-tagged oligos C-2 NF to C-5 NF (with 2-5 repeats of 5’-CCCCGG-3’ sequence) and a double-tagged C-rich Myc oligo (cMyc C FC) were treated without or with MEI in GMH1 buffer. **J**. Cy5-tagged oligos C-2 NF to C-5 NF (with 2-5 repeats of 5’-CCCCGG-3’ sequence) and a FAM-tagged G-2 oligo treated without or with MEI in GMH2 buffer.

Further, we explored MEI cleavage limitations as regards the absence or presence of flanking sequences outside of a G4-forming sequence. We used several oligos containing varying numbers (1-6 and 14) of 5’-GGGGCC-3’ repeats, but without any flanking sequences (G-1 NF – G-6 NF and G-14 NF). As shown in **Fig. 2 E and Fig. S1 A-B**, MEI did not cleave oligos containing 1 or 2 repeats but cleaved three oligos containing 3, 4 and 5 hexanucleotide repeats in GMH1 and NEBuffer r2.1. We then treated two oligos containing 6 repeats (G-6 NF) and 14 repeats (G-14 NF) of 5’-GGGGCC-3’ without any flanking sequence with MEI in three different buffers (GMH1, GMH2 and GMH3 buffers). The oligo containing 6 repeats without flanking sequences (G-6 NF) generated several bands, but the band with the smallest size was the major cleavage product (**Fig. 2 F-H**). The oligo containing 14 repeats without any flanking sequence (G-14 NF) generated several prominent bands (**Fig. 2 F-H**). These results suggest that MEI can cleave an oligo if it contains 3 or more repeats of 5’-GGGGCC-3’ sequence without a flanking sequence. The enzyme can cleave an oligo containing 1 and 2 repeats of 5’-GGGGCC-3’ sequence if the repeats are flanked by DNA sequences to provide space and site(s) for cleavage.

Since MEI cleaved G4-forming oligos containing repeats of 5’-GGGGCC-3’, we wanted to test if the enzymatic activity is specific to G4-forming oligos or if it can cleave oligos that form other structures such as the i-motif as is generated by the complementary sequence 5’-CCCCGG-3’. We designed and treated four different oligos containing 2-5 hexanucleotide repeats of 5’-CCCCGG-3’ sequence with MEI in two different buffers (GMH1 and GMH2 buffers). The oligos (**Supplementary Materials; Table S1**) do not contain flanking sequences, and they were tagged with Cy5 dye that emits bright red fluorescence. Gel analysis showed that MEI did not cleave any of the polycytosine-rich DNA oligos (**Fig. 2 I-J**). The above results on oligos containing 5’-GGGGCC-3’ and 5’-CCCCGG-3’ sequences show that MEI cleaves oligos containing polyguanine-rich hexanucleotide repeats of the *C9orf72* that form G4 but does not cleave oligos containing polycytosine-rich repeats. The results also showed that MEI requires flanking sequences outside of G4 forming sequences for cleavage. Finally, the results also showed that MEI cleavage of G4-forming sequence of *C9orf72* generates banding patterns, a diagnostic feature of the repeat numbers.

### Mismatch Endonuclease I specifically cleaves G4-forming sequences present at oncogene promoter-proximal regions and telomeres

Since MEI recognized and cleaved all G4-forming oligos containing 5’-GGGGCC-3’ repeats of *C9orf72*, but not i-motif-forming oligos containing 5’-CCCCGG-3’ repeats, we inquired if the enzyme would site/structure-specifically recognize and cleave other G4-forming oligos. Using CD spectroscopic analysis of secondary structures formed by polyguanine-rich DNA and their complementary polycytosine-rich DNA sequences from the promoter-proximal regions of several oncogenes including cMyc, BCl2, EGFR, VEGF, and hRAS, we have recently analyzed the effects of pH on G4 and i-motif formation ^11^. To determine the effects of MEI on G4s formed by these sequences, we designed oligos that contained (**a**) polyguanine-rich (G) and polycytosine-rich (C) sequences of cMyc, BCl2, EGFR, VEGF, and hRAS, (**b**) 8-10 nucleotides flanking sequences on either sides of the G4- and i-motif-forming sequences,(**c**) a FAM tag on their 5’ ends, and (**d**) a Cy5 tag on their 3’ ends (double-tagged). In addition, two cMyc G oligos (that form G4s) were also designed, one with a 5’ FAM tag alone (cMyc G 5’F) and a second oligo with a 3’ FAM tag alone (cMyc G 3’F) (**Supplementary Materials, Table S1**). We first conducted CD analysis of an untagged cMyc G oligo (cMyc G) and the two FAM-tagged cMyc G oligos (cMyc G 5’F and cMyc G 3’F). The untagged cMycG oligo displayed a positive peak at 265 nm and a negative peak at 240 nm typical for cMyc G4 in sodium cacodylate buffer containing 100 mM KCl at pH 7.4 (**Supplementary Materials, Table S1**). CD spectroscopic analysis of the 5’-FAM-tagged cMycG (cMyc G 5’) and the 3’-FAM-tagged cMycG (cMyc G 3’F) each showed a positive peak at ∼265 nm and a negative peak at ∼240 nm as the untagged cMycG oligo (**Supplementary Materials, Fig S1**) suggesting that tagging did not affect G4 formation.

We treated all dual-tagged and single-tagged G4- and i-motif-forming oligos containing sequences from the oncogene promoter-proximal regions with MEI enzyme and analyzed them in denaturing gels (**Fig. 2 I and 3 A-F**). A dual-tagged i-motif-forming polycytosine-rich oligo of cMycC was not cleaved by MEI (**Fig. 2 I**). However, MEI enzyme cleaved all four dual-tagged polyguanine-rich, G4-forming oligos containing sequences from the promoter-proximal regions of cMycG, BCL2G, EGFRG, and hRASG in GMH1, GMH2, GMH3 and NEBuffer r2.1 buffers and showed multiple red and green fluorescent bands (**Fig. 3 A-D, cMycG 8FC, BCL2G 10FC, EGFRG 10FC**, and **hRASG 1-FC**). In addition, MEI treatment of a 5’-FAM-tagged G4-forming cMyc oligo (cMyc G 5’F) showed a single cleavage product in GMH1 buffer (**Fig. 3 A cMycG 10F**). Further, the enzyme also cleaved a 5’-FAM-tagged oligo containing the telomeric sequence (Tel G 5F) generating a green, fluorescent band (**Fig. 3 A Tel G 5F and Fig. 3 C TelG 5F**). Since all oligos had a FAM tag on their 5’ ends each, we wanted to ascertain that the FAM-tag did not influence the enzymatic cleavage process. We tagged an oligo containing a G4-forming polyguanine-rich sequence associated with fragile X syndrome (Fragile X G) with Cy5 and treated it with MEI in GMH buffer1. The enzyme cleaved the oligo to generate multiple red fluorescent bands (**Fig. 3 E, Fragile X G**). To determine the specificity of MEI to polyguanine-rich, but not polycytosine-rich, DNA, we designed Cy5-tagged oligos containing i-motif-forming polycytosine-rich sequences from the promoter-proximal regions of cMyc, BCl2, EGFR and hRAS genes with varying number of flanking sequences (**Supplementary Materials, Table S1**); these sequences are complementary to the polyguanine-rich DNAs used above. The sequences can form i-motifs in buffers at low pH (pH 5.5 -6.5)^11^. As shown in **Fig 3 E**, MEI did not cleave any of the oligos containing polycytosine-rich DNA sequences. We also designed four FAM-tagged oligos that do not form G4s or i-motifs (**Supplementary Materials, Table S1**) and evaluated them for cleavage by MEI. As shown in **Fig. 3 F**, MEI did not cleave any of the four oligos, namely ODN1, ODN2, ODN4, and ODN5. All results in **Fig. 3** show that MEI cleaved only G4-forming polyguanine-rich sequences present at the promoter-proximal regions of several oncogenes and telomeres but did not cleave i-motif-forming sequences and other non-G4-forming sequences.

**Fig. 3.**
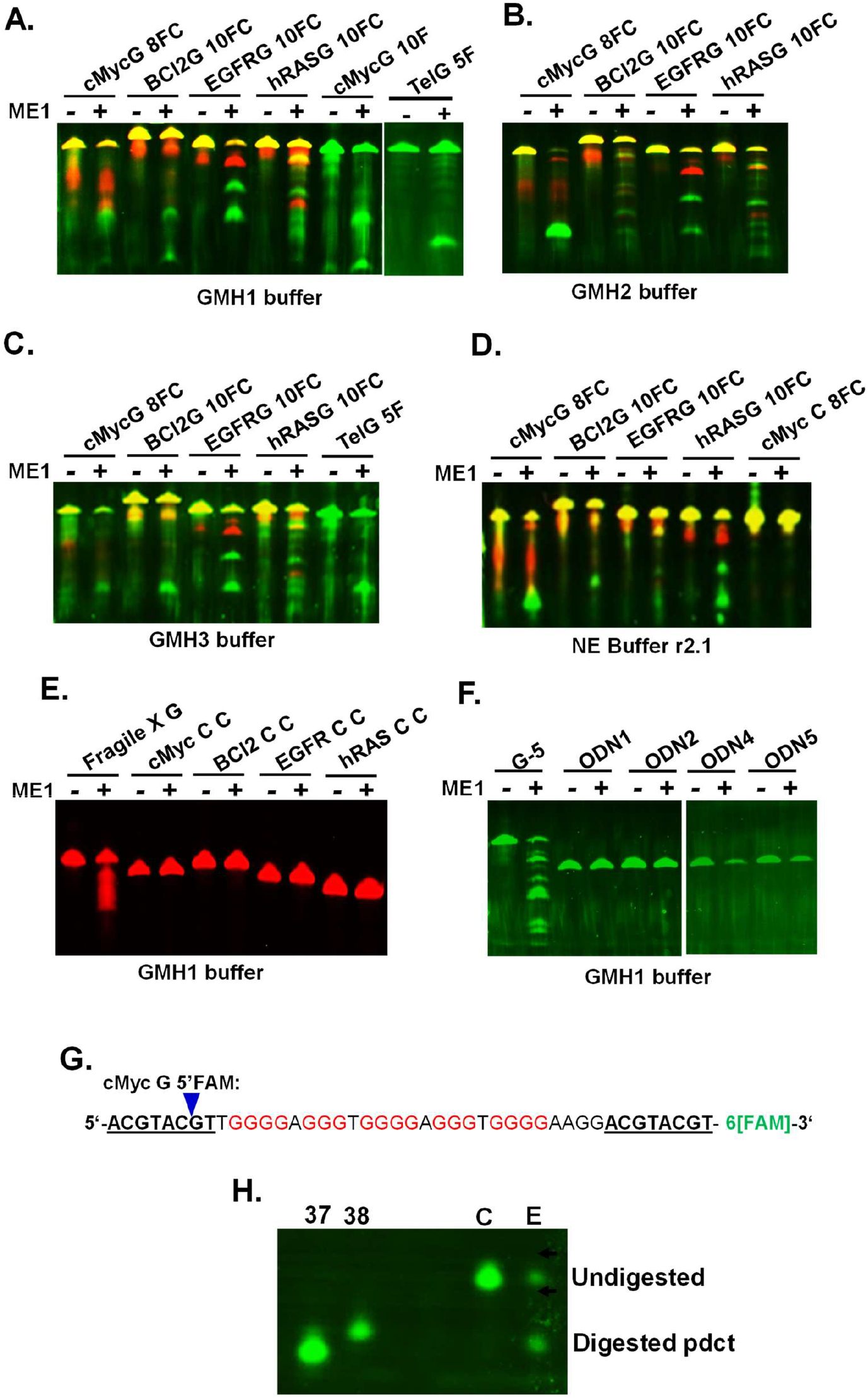
7M urea-20% polyacrylamide gels show that Mismatch Endonuclease I (MEI) cleaves G4-forming, but not i-motif-forming, oligos containing sequences of oncogene-promoter proximal region and sequences of telomeres in a site-specific manner. Yellow color is due to the presence of FAM-tagged and Cy5-tagged oligos of the same size whereas green color is due to FAM-tagged DNA and red color is due to Cy5-tagged DNA fragments. MEI (-) and (+) denote lanes containing DNA without and with MEI. **A-D**. The gels show MEI digestion of G4-forming dual tagged oligos from the promoter-proximal regions of cMyc (**cMycG 8FC**), BCL2 (**BCl2G 10FC**), EGFR (**EGFRG 10FC**), and hRAS (**hRASG 10FC**) in GMH1, GMH2, GMH3 and NEBuffer r2.1 buffers. In addition, (**A**) shows FAM-tagged oligos from cMyc promoter region (**cMycG 10F**), (**A and C**) show a FAM-tagged oligo from telomere (**TelG 5F**), and (**D**) shows a dual tagged cMyc oligo (**cMyc C 8FC**). **E**. MEI digestion of Cy5-tagged G-rich DNA of fragile X (**Fragile X G**), and Cy5-tagged C-rich DNAs of cMyc (**cMyc C C**), BCL2 (**BCL2 C C**), EGFR (**EGFR C C**) and hRAS (**hRAS C C**). **F**. MEI digestion of FAM5-tagged G-rich DNA of G-5, ODN1, ODN2, ODN4 and ODN5. **G**. Sequence of the cMyc DNA oligo used to map the MEI cleavage site. The letters underlined and in bold are 8 additional nucleotides added to each end of traditionally used 27-base-long G4-forming cMyc oligo. The letters colored red are guanine sequences involved in the G4 formation. The downward facing arrowhead points to the MEI cleavage site. **H**. Mapping of MEI cleavage site. The oligos numbered 37 and 38 are FAM-tagged 37- and 38-nucleotide long oligos used as molecular markers. **Lane ‘C’**: oligo without MEI treatment (negative control), and **Lane ‘E’**: digestion by MEI (enzyme). The two green fluorescent bands in the lane are undigsted DNA (top) and MEI-cleaved band (bottom; 37 nucleotides-long).

We have examined the exact site of MEI cleavage in a G4-forming sequence. The 43-nucleotide long 3’-FAM-tagged oligo (cMyc G 3F) contained the 27-nucleotide long G4-forming sequence of the cMyc promoter region with 8 additional nucleotides on each end (**Fig. 3 G**). The oligo was treated with MEI, and the products were fractionated in a denaturing PAGE gel along with several oligos of known size as molecular weight standards. The MEI digestion resulted in the generation of a FAM-tagged 37-nucleotide long product (**Fig. 3 I**); the 6 nucleotide long untagged fragment is not visible in the gel as it does not have a tag. The generation of the 37-nucleotide long FAM-tagged product suggests that the enzyme cleaved 3 nucleotides upstream of the first guanine of the G4 in the 43-nucleotide long cMyc oligo.

### HaeIII restriction enzyme recognizes and cleaves the hexanucleotide repeats of *C9orf72* gene generating specific patterns

The polyguanine-rich DNA containing hexanucleotide repeats of 5’-GGGGCC-3’ forms G4s at around neutral pH (e.g., pH 7.4) in the presence of monovalent cations such as potassium and sodium.. To demonstrate HaeIII restriction endonuclease enzyme’s ability to assess the number of hexanucleotide repeats in conditions favoring both the formation of G4s and GGCC/CCGG base pairs, we treated various oligos containing 1-5 repeats of 5’-GGGGCC-3’ with flanking sequences, oligos containing 1-5 repeats without any flanking sequences, and oligos containing 6 and 14 repeats without any flanking sequences (**Supplementary Materials, Table S1** and oligos used in **Fig. 2**) with HaeIII restriction endonuclease. As shown in **Fig. 4 A-B**, the enzyme cleaved all oligos containing 1-5 repeats of 5’-GGGGCC-3’ and flanking sequences (G-1 – G-5) in both GMH1 buffer and NEBuffer r2.1 and generated specific banding patterns. We then tested oligos containing 1-6 repeats of 5’-GGGGCC-3’ (G-1 NF -G6 NF) as well as an oligo containing 14 repeats of 5’-GGGGCC-3’ (G-14 NF) for MEI cleavage; all these oligos lacked other flanking sequences. While the enzyme did not cleave the G-1 NF oligo containing one repeat, it cleaved a small fraction of the G-2 NF oligo (**Fig. 4 C-D**) and a significant fraction of the remaining oligos, namely G-3 NF to G-14 NF, generating multiple bands (**Fig. 4 C-D**). However, the enzyme did not cleave G4-forming oligos containing sequences from the proximal-promoter regions cMyc, BCl2, EGFR and hRAS oncogenes or i-motif-forming sequences (**Fig. 4 D-F**).

**Fig. 4.**
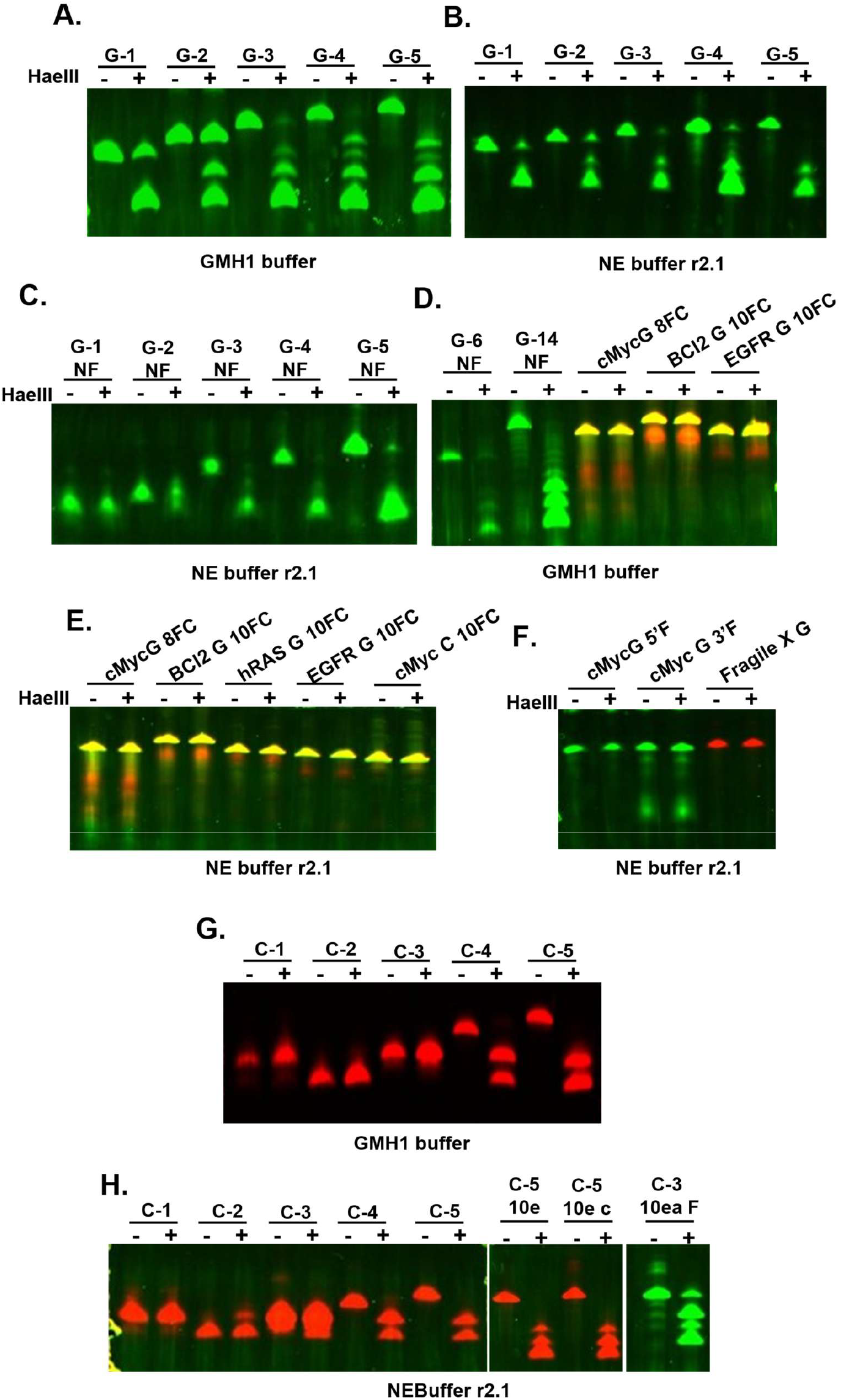
7M urea-20% polyacrylamide gels show that HaeIII enzyme cleaves G4-forming DNA oligos containing repeats of *C9orf72* gene. A-H. (-) denotes without enzyme and (+) denotes with HaeIII enzyme. **A**. HaeIII digestion of FAM-tagged G-1 to G-5 oligos containing flanking sequences in GMH1 buffer. **B**. HaeIII digestion of FAM-tagged G-1 to G-5 oligos containing flanking sequences in NEBuffer r2.1. **C**. HaeIII digestion of FAM-tagged G-1 NF to G-5 NF oligos without any flanking sequences in NEBuffer r2.1. **D**. HaeIII digestion of oligos containing FAM-tagged G-6 NF and FAM-tagged G-14 NF, and dual tagged G4-forming cMyc, BCL2 and EGFR (cMycG 8FC, BCL2G 10Fc and EGFRG 10FC) oligos in GMH1 buffer. **E**. HaeIII digestion of oligos containing dual tagged G4-forming cMyc, BCL2, hRAS and EGFR (cMycG 8FC, BCL2G 10FC, hRASG 10FC, and EGFRG 10FC) oligos and i-motif-forming cMyc oligos (cMycC 10FC) in NEBuffer r2.1. **F**. HaeIII digestion of cMyc G 5F, cMyc G 3’G and Fragile X ggc C oligos in NEBuffer r2.1. **G**. HaeIII digestion of C-1 NF to C-5 NF oligos in GMH1 buffer. **H**. HaeIII digestion of oligos C-1 NF– G-5 NF, C-5 10e, C-5 10ec, and C-3 10ea F in NEBuffer r2.1.

Our results show that while MEI cleaved all G4-forming oligos from *C9orf72*, oncogene promoter-proximal regions, and telomeres, HaeIII cleaved GGCC/CCGG base-pair forming oligos from *C9orf72*, but not the oligos from the oncogene-promoter proximal regions. Based on our prior finding that MEI does not cleave oligos containing 5’-CCCCGG-3’ sequence (**Fig. 2 I-J**)we treated oligos containing 1-5 repeats of 5’-CCCCGG-3’ with HaeIII and observed that the enzyme cleaved oligos containing 4 and 5 repeats in GMH1 buffer and NEBuffer r2.1 (**Fig. 4 G-H**), consistent with their potential to form GGCC/CCGG base pairs. When we added additional flanking sequences to an oligo containing 5 repeats of 5’-CCCCGG-3’, HaeIII digestions resulted in an increased number of fragments (**Fig. 4 H**).

### MEI and HaeIII together cleave G4-forming repeats of *C9orf72* generating specific patterns

Since both MEI and HaeIII cleaved the G4-forming oligos containing 5’-GGGGCC-3’ repats, we tested these substrates for simultaneous digestion with both enzymes. We treated oligos containing 1-5 repeats with flanking sequences with MEI, HaeIII, and both MEI and HaeIII with one enzyme added after incubation with the other or simultaneously. As shown in **Fig. 5 A**, each of MEI and HaeIII cleaved every oligo to generate an individual banding pattern of the latter. However, when the oligo contained 1 or 2 repeats, double digestion with both MEI and HaeIII generated a pattern identical to that generated by MEI digestion. When the oligo contained 3-5 repeats, the banding pattern of the double digestion looked like that generated by the addition of the first enzyme.

**Fig. 5.**
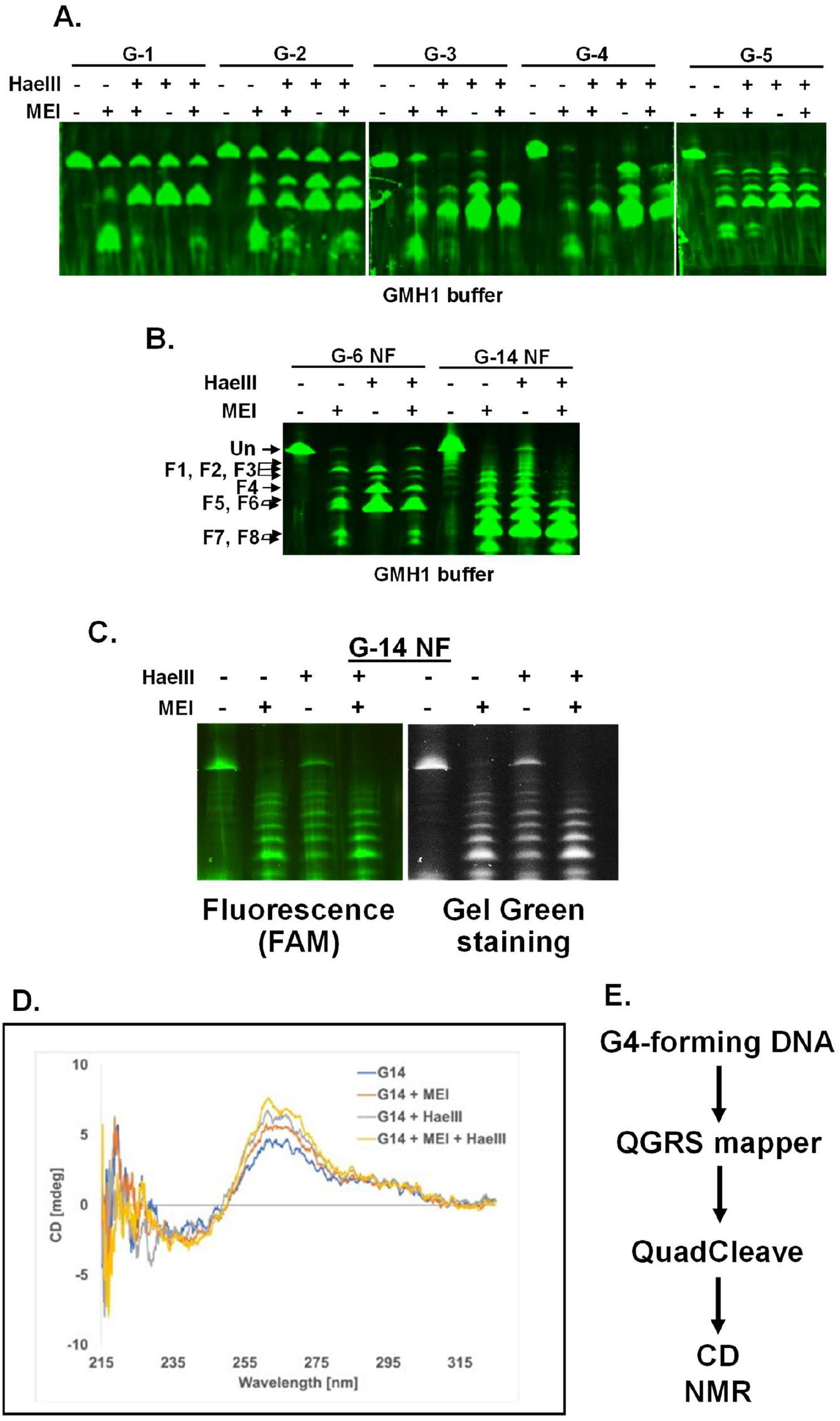
7M urea-20% polyacrylamide gels show double-digestion of G4-forming oligos of *C9orf72* by Mismatch Endonuclease I and HaeIII enzymes generating specific pattern of DNA bands. **A**. MEI and HaeIII double-digestion of FAM-tagged oligos G-1 – G-5 in GMH1 buffer. **B**. MEI and HaeIII digestion of FAM-tagged G-6 NF and G-14 NF oligos. **C**. Double-digestion of a mixture of untagged and FAM-tagged G-14 NF oligo by MEI and HaeIII enzymes for CD spectroscopic analysis. **D**. CD spectroscopic analysis of G4 formation by a mixture of untagged and tagged G-14 NF oligo after single- or double-digestion by MEI and HaeIII. **E**. Flow chart showing methods for G4 analysis of an oligo under test.

When an oligo contained 6 or 14 repeats of 5’-GGGGCC-3’ without flanking sequences and was incubated with MEI, HaeIII, or MEI+HaeIII, the single digests generated their individual patterns whereas the double digestion with MEI and HaeIII generated a banding pattern of MEI (**Fig. 5 B-C**). We conducted a CD spectroscopic analysis of single digests and double-digests of G-14 NF oligo (that contained 14 repeats of 5’-GGGGCC-3’ without flanking sequences) and observed that the CD values increased marginally upon cleavage by the two enzymes. All these results demonstrate that *C9orf72* oligos are amenable to enzymatic digestions by MEI and HaeIII.

## Discussion

The numerous polyguanine-rich sequences present in the human genome can form G4 secondary structure at different times in the cell cycle, which can regulate various cellular transactions on DNA including transcription and replication. The G4s can cause DNA damage which can lead to genetic diseases such as cancer, ALS, FTD and ALS-FTD. All other organisms also contain numerous potential G4-forming sequences that can regulate various cellular processes and genome integrity. Thus, identification and analysis of a G4 formed by a DNA sequence will help understand the biological functions of the sequence. Inexpensive methods for the identification and confirmation of an individual G4 *in vitro* is required. QGRS mapper can quickly suggest if a DNA sequence can form a G4 structure, however, it requires a method to validate the prediction of a G4 formation and determine the number of G4s formed by a DNA fragment. QuadCleave is the first enzymatic technique to recognize and site-specifically cleave G4-forming DNA. Various helicases and nucleases exist in the cell to resolve G4s^40–44^; however, no helicase or nuclease has yet found universal application for *in vitro* or *in vivo* analysis of G4s. Several biophysical techniques are used to study and analyze G4s. CD spectroscopy is one of the most common methods to identify and study G4s^11^; however, the equipment is relatively expensive. Similarly, NMR^45^ and X ray crystallography^46,47^ have been used to study G4s, but are more expensive than a CD spectroscope. The enzymatic analysis of G4-forming ssDNA oligos by ME1 is specific, commercially available, and is inexpensive. Using our approach, MEI recognized and cleaved all G4-forming substrates tested so far and did not cleave other types of DNA substrates including several i-motif-forming substrates and linear DNA. QuadCleave can be conducted easily to assess a potential G4-forming DNA.

MEI digestion of G4-forming sequences of *C9orf72* hexanucleotide containing flanking sequences generates a pattern that displays unique characteristics of the number of hexanucleotide repeats. Therefore, the current finding holds promise in the identification and analysis of the copy numbers of the G4-forming repeats in the test DNA of *C9orf72*. Experiments are underway to apply QuadCleave for the analysis of hexanucleotide copy numbers in chromosomal DNA preparations *in vitro* and *in vivo*.

The finding that HaeIII cleaves both hexanucleotide repeats 5’-GGGGCC-3’ and 5’-CCGGGG-3’ of *C9orf72* shows that oligos containing the *C9orf72* hexanucleotides can form 5GG/CC’ base pairs. The double-digestion of the hexanucleotide repeats by the 2-enzyme system of MEI and HaeIII shows that both G4 structure and GG/CC base pairs coexist in the G4-forming 5’-GGGGCC-3’ repeats of *C9orf72* and can be used for the analysis of the hexanucleotide repeats *in vitro*. QuadCleave also holds promise for *in vivo* G4 analysis.

MEI enzyme not only recognizes and cleaves G:G mismatch but also can cleave G:T and T:T mismatches in a double-stranded DNA. It also cleaves T:I, G:I and G:U mismatches in DNA as well as T:G and T:U mismatches in DNA:RNA hybrid. The enzyme can also nick the thymine strand of T:C mismatches in DNA. Various organisms in the groups of archaea and *Mycobacteria* produce endonucleases such as EndoMS or NucS that recognize and cleave G:G, G:T and T:T mismatches by mismatch repair (MMR) system have been identified^48–52^. Experiments are underway to explore the potential that EndoMS/NucS will recognize and site-specifically cleave G4s as MEI does. In addition, if the mismatches discussed above are present in a G4-forming DNA, MEI and EndoMS/NucS may cleave such mismatches in addition to cleaving the G4. Therefore, a mutant MEI or EndoMS/NucS enzyme that specifically cleaves G:G mismatch, but not the other mismatches, would also be an ideal enzyme for G4 analysis; such studies are underway. Further, it is also possible that a G4-forming DNA sequence contains any of the above-mentioned mismatches (G:G, G:T etc). In such substrates, MEI will cleave 2 and 3 nucleotides upstream of the mismatch and the G4 structure, complicating the analysis and conclusions. Since MEI cleaves the DNA outside of the G4 structure(s), CD and/other analysis can be used in combination as a follow-up to the analysis of MEI cleavage products to circumvent such issues.

QuadCleave can be used in genome-wide preparation, identification, and analysis of G4s from a DNA library, and experiments are underway on genome-wide experiments. In conclusion, QuadCleave is an enzymatic method to identify and analyze G4s formed by a DNA sequence from any organism and in combination with HaeIII enzyme, QuadCleave can be used in the biochemical analysis of G4 formed by the hexanucleotide repeats in intron 1 of *C9orf72* associated with ALS and FTD.

## Methods

### Oligonucleotides

Oligonucleotides (oligos) used in the study are listed in Table S1. All oligos were obtained from EMD Millipore with a few exceptions from IDT.

### Mismatch Endonuclease I digestion

An MEI digestion reaction (25 µl) contained 8 pmoles of DNA and without or with 80 units of MEI enzyme (New England Biolabs) in one of the following (G4-MEI-HaeIII or GMH) buffers - *<u>GMH buffer 1</u>*: 10 mM Na.Cacodylate (pH 7.4), 100 mM KCl, 10 mM MgCl_2_, 100 µg/ml acetylated BSA; *<u>GMH buffer 2</u>*: 10 mM Na.Cacodylate (pH 7.4), 50 mM NaCl, 10 mM MgCl_2_, 100 µg/ml acetylated BSA. *<u>GMH buffer 3</u>*: 10 mM Tris.HCl (pH 7.5), 50 mM NaCl, 10 mM MgCl_2_, 100 µg/ml acetylated BSA. *<u>GMH buffer 4</u>*: 10 mM sodium cacodylate (pH 7.4), 50 mM NaCl, 10 mM KCl, 10 mM MgCl_2_, 100 µg/ml Recombinant Albumin. The reaction mix was incubated at 37°C for 1-2 hours or 16-20 hours.

### HaeIII restriction endonuclease digestion

A HaeIII digestion reaction (25 µl) contained 8 pmols of DNA and without or with 10 units of HaeIII (New England Biolabs) in one of the following buffers - *<u>GMH buffer 1</u>*: Na.Cacodylate (pH 7.4), 100 mM KCl, 10 mM MgCl2, 100 µg/ml acetylated BSA; *<u>GMH buffer 2</u>*: 10 mM Tris.HCl (pH 7.5), 50 mM NaCl, 10 mM MgCl2, 100 µg/ml acetylated BSA. The reaction mix was incubated at 37°C for 1-2 hours or 16-20 hours.

### Urea polyacrylamide gel electrophoresis (urea-PAGE)

The DNA samples were fractionated in a urea-PAGE containing 7 M urea and 10%-20% polyacrylamide. The gels were run at 170 volts. The fluorescence-tagged DNA samples were analyzed by iBright 1500 gel documentation unit in the universal channel for FAM and Cy5.

### Circular dichroism (CD) spectroscopy

CD spectroscopy was carried out essentially as described ^11^ with modification in buffers. The buffer used for G4 was either GMH1 buffer or 10 mM Na.cacodylate + 100 mM KCl ^11^.

## Acknowledgments

BKM was funded by VCOM’s REAP grants 1032453 and 1302559.

## Additional information

**Correspondence** and requests for materials should be addressed to.

---

**Fig. S1.**
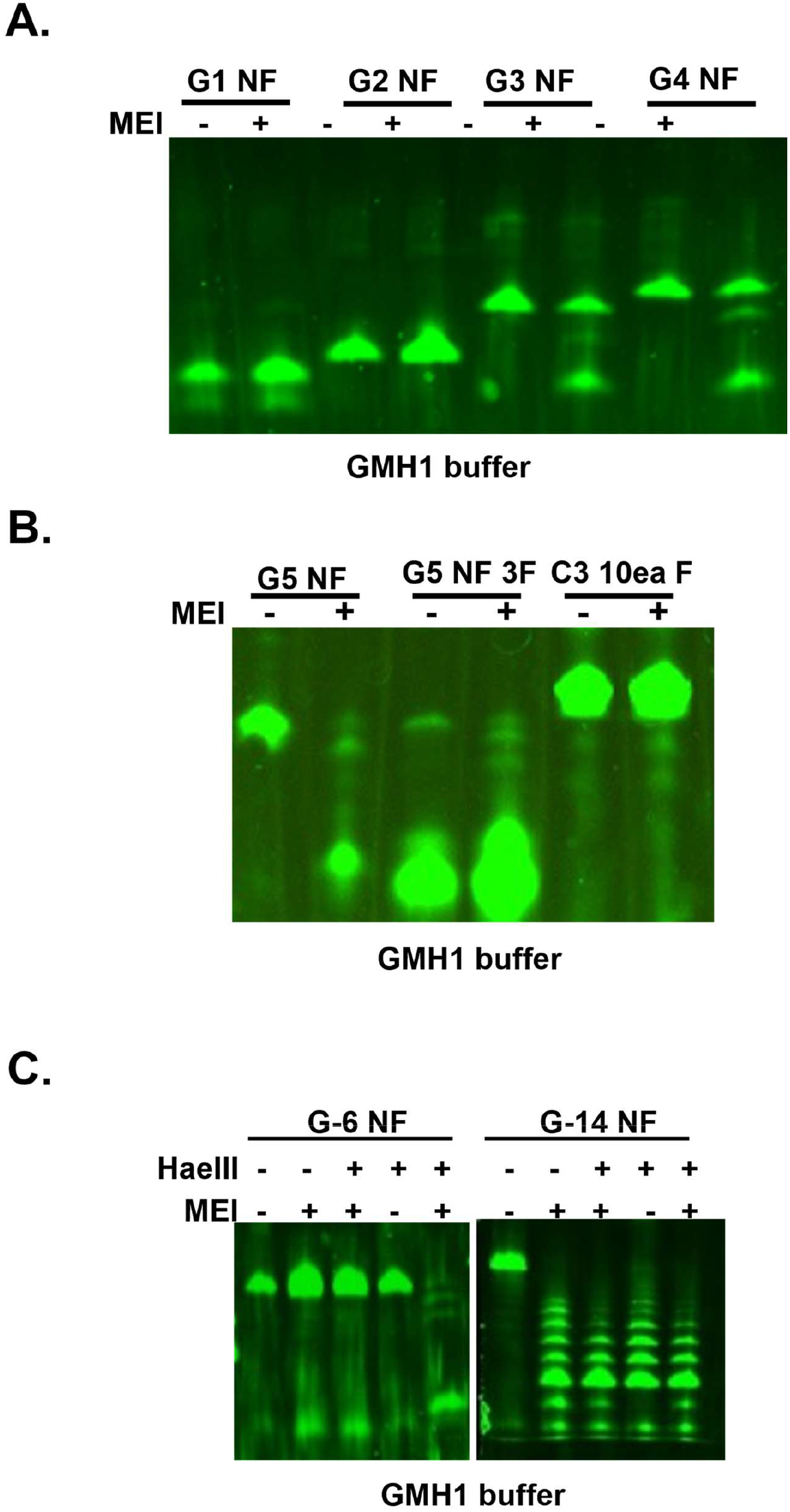
7M urea-20% polyacrylamide gels show MEI- and HaeIII-digestion of oligos containing various G4 forming repeats of 5’-GGGGCC-3’ without flanking sequences. **A**. MEI cleavage of G4-forming oligos G-1 NF – G-4 NF in GMH1 buffer. **B**. MEI cleavage of G4-forming oligos 5’-FAM-tagged G-5 NF and 3’-FAM-tagged G-5 NF containing 5’-GGGGCC-3’ sequence, and i-motif-forming 5’-FAM-tagged 5’-CCCCGG-3’ oligo (C-3 10ea F) in GMH1 buffer. **C**. Double-digestion of 5’-FAM-tagged G4-forming oligos G-6 NF and G-14 NF by MEI and HaeIII in GMH1 buffer.

**Fig. S2.**
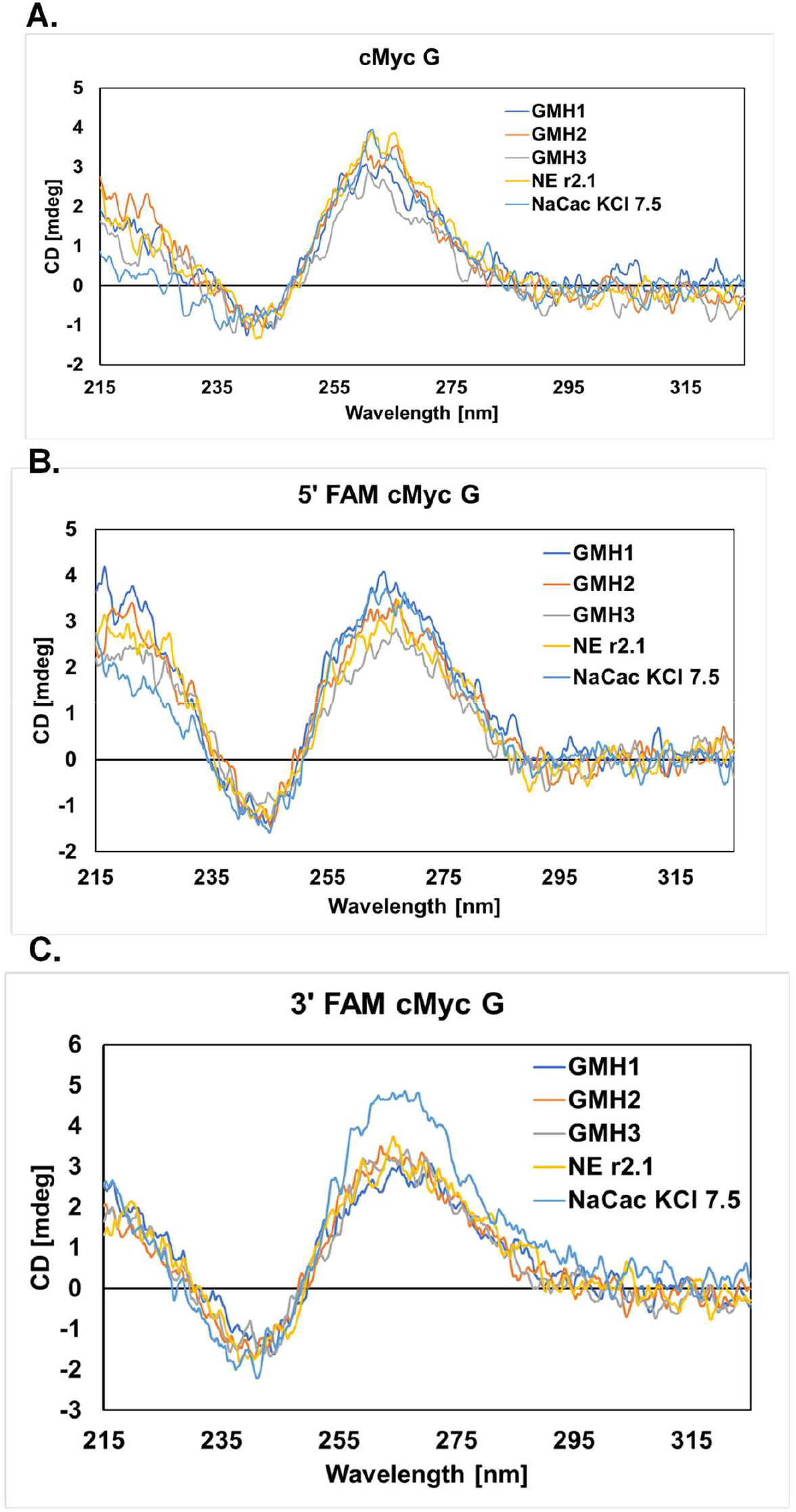
CD spectroscopic analysis of G4-forming cMyc G oligos in five different buffers, sodium cacodylate/KCl (pH 7.4), GMH 1, GMH2, GMH3 and NEBuffer r2.1. **A**. CD analysis of an untagged cMyc G oligo containing 27 nucleotides. **B**. CD analysis of a 5’-FAM-tagged cMycG oligo with 43 nucleotides (27 nucleotides as in A with 8 additional flanking nucleotides on each end). **C**. CD analysis of a 3’-FAM-tagged cMycG oligo with 43 nucleotides (27 nucleotides as in A with 8 additional flanking nucleotides on each end).

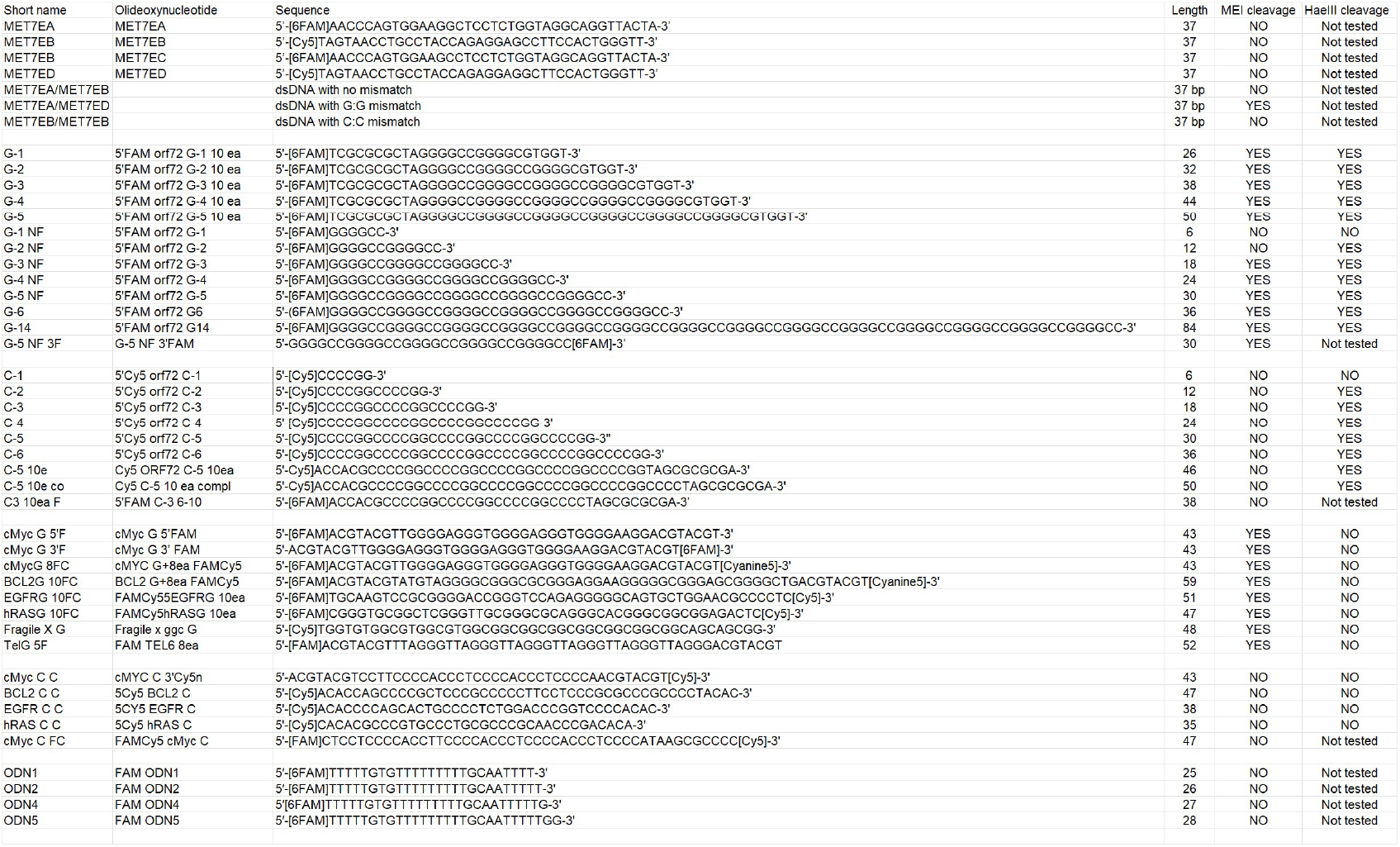

